# Chemokine receptor activity is differentially regulated by membrane cholesterol

**DOI:** 10.64898/2026.07.20.738555

**Authors:** Fernando Salgado-Polo, Jose Fernández-González, Carolina Ferrera-Mena, Suresh Subedi, Sriram Tiruvadi-Krishnan, Philip Ben Rainsford, Rajesh Regmi, Rajan Lamichhane, Martin Gustavsson

**Affiliations:** Department of Biomedical Sciences, Faculty of Health and Medical Sciences, University of Copenhagen, Blegdamsvej 3B, 2200 Copenhagen, Denmark; Department of Biochemistry & Cellular and Molecular Biology, University of Tennessee, Knoxville, Tennessee, USA

**Keywords:** G protein-coupled receptor, Receptor activation, Oxysterol, Nanodisc, Cyclodextrin, CXCR4, Single-molecule FRET

## Abstract

Cholesterol is a key membrane component that regulates G protein-coupled receptor (GPCR) function, yet its molecular mechanisms remain unclear. Here, we combine chemical extraction of membrane sterols with functional signaling assays and single-molecule fluorescence resonance energy transfer (smFRET) to define how cholesterol controls activation of chemokine receptors. Reduction of membrane cholesterol in mammalian cells selectively decreased constitutive and agonist-induced signaling across CXCR1, CXCR2, CXCR4, while it activated ACKR3, and did not affect CXCR3, revealing receptor-specific dependence on membrane sterols. Mechanistically, cholesterol regulation partly required the conserved class A GPCR residue Trp^4.50^ and shifted agonist-bound CXCR4 toward active conformational states, providing a molecular explanation for its functional effects. In contrast, replenishment with oxidized cholesterol species failed to restore receptor activity, distinguishing cholesterol from oxysterols as modulators of receptor activation. Our findings identify cholesterol as an allosteric regulator of chemokine receptors and suggest that oxysterols may reshape inflammatory signaling by selectively modulating GPCR activity.

## Main

Cell membranes are dynamic structures, whose charge, thickness and rigidity strongly influence protein structure and function^1^. Sterols such as cholesterol are key membrane components that reduce permeability, modify membrane organization, and influence protein dynamics and receptor signaling. In particular, G protein-coupled receptors (GPCRs) are regulated by membrane composition, affecting trafficking, oligomerization and signaling^2^. For example, cholesterol acts as an allosteric modulator of the adenosine A_2A_ receptor, shifting conformational equilibria and enhancing agonist-dependent activation through direct receptor–sterol interactions^3,4^.

Similar lipid-dependent modulation has been observed in chemokine receptors^5–7^, a family of GPCRs that mediate immune regulation, and inflammatory processes upon binding small (8–12 kDa) proteins known as chemokines^8^. Depending on their ligand preference, chemokine receptors are classified in CC, CXC, CX3C and C subfamilies^9^. Among the CXC chemokine receptors, CXCR4 has been well-characterized for its essential roles in immune cell migration and embryonic development upon binding to the chemokine CXCL12. In addition to CXCR4, CXCL12 also binds to CXCR4, ACKR3, and ACKR5: while all three receptors can recruit β-arrestin upon activation, only CXCR4 triggers G protein-driven signaling leading to immune cell migration^10^.

The cell signaling functions of CXCR4 have been repeatedly reported to be regulated by cholesterol^11,12^. For instance, cholesterol-rich membrane microdomains regulate CXCR4 surface localization and CXCL12 binding, while cyclodextrin-mediated cholesterol depletion leads to reduced chemotaxis and receptor stability^13,14^. In pathological contexts, oxidized cholesterol species have been reported to impair CXCR4 signaling in senescent endothelial and immune cells, linking lipid oxidation to aging-associated immune dysfunction^15–17^. Cholesterol has also been implicated in receptor oligomerization^18,19^, such as the homo- and heterodimer formation among CXCR4 and CCR5, which partition into cholesterol-rich microdomains during T-cell activation, facilitating HIV-related cell-cell interactions^20,21^. However, it remains unresolved whether cholesterol mainly alters receptor expression and ligand engagement or whether it influences CXCR4 conformational states and signaling.

Structural approaches have further identified and examined the association of chemokine receptors with sterols^22–25^. Direct sterol binding has been predicted to determine the structure and stability of 44% of class A GPCRs^26^ through the cholesterol consensus motif (CCM) (Arg/Lys^4.39–4.43^−Trp/Tyr^4.50^−Ile/Val/Leu^4.46^−Phe/Tyr^2.41^), where Trp^4.50^ (residue numbering follows the Ballesteros/Weinstein scheme) is almost universally conserved (94%) and interacts with sterols in many structures^26,27^. Addressing how cholesterol influences chemokine receptor activity at a molecular level has become technically feasible through membrane reconstitution using nanodiscs. Nanodiscs are soluble, nanoscale lipid bilayers that mimic biological membranes and enable biophysical characterization in chemically defined lipid environments^28,29^. Moreover, single-molecule fluorescence resonance energy transfer (smFRET) has enabled real-time visualization of the conformational dynamics of CXCR4 and ACKR3 upon ligand binding^30^.

Here, we investigated how membrane cholesterol regulates signaling and dynamics of CXC chemokine receptors. Reduction of cholesterol levels in mammalian cells selectively impaired CXCR4-mediated constitutive and ligand-induced G protein signaling and β-arrestin recruitment, but only partially affected CXCR1 and CXCR2 and did not compromise CXCR3 or ACKR3 activation. Mechanistically, this regulation partly involved the conserved GPCR residue Trp^4.50^, and smFRET showed that cholesterol promotes transitions of agonist-bound CXCR4 toward specific active receptor states. In contrast, oxysterols failed to restore activity. Overall, our findings determine how cholesterol composition regulates receptor signaling, suggesting how altered cholesterol chemistry may contribute to inflammatory disease pathology.

## Results

### Cholesterol selectively modulates β-arrestin recruitment to CXCR4

To understand the role of cholesterol in chemokine receptor activity, we extracted it from HEK293A cells using hydroxypropyl–β–cyclodextrin (HPβC) (**Fig.1a**). HPβC has been previously reported to scan the plasma membrane, where it can insert to embed cholesterol molecules, thereby extracting cholesterol^31^. We first optimized the amount of HPβC needed to extract detectable levels of cholesterol (**Fig.S1a-c**), which neither reduced cell membrane rigidity nor affected cell viability (**Fig.S1d-e**).

We next determined the effect of cholesterol depletion on the recruitment of β-arrestin-2 to CXCR4. For that, we utilized *Renilla* luciferase (Rluc)-tagged CXCR4 and measured bioluminescence resonance energy transfer (BRET) to venus-tagged β-arrestin-2 upon stimulation with its natural agonist, CXCL12 (**Fig.1b**). Our results show that the efficacy (or E_max_) for CXCL12 to enact β-arrestin coupling was reduced by HPβC in a concentration-dependent manner without significant changes in ligand potency (or EC_50_) or basal activity (**Fig.1c-d, S1f**), which suggests that cholesterol may be important for CXCR4 activation without affecting CXCL12 binding.

To test if the effects of cholesterol are reversible, we restored cholesterol in the cell membrane using cholesterol-loaded HPβC (**Fig.1a**). This treatment effectively doubled the amount of cholesterol, which significantly increased cell rigidity without changes in cell viability (**Fig.S1c-d**). Importantly, cholesterol restoration rescued the efficacy of CXCL12 to trigger β-arrestin-2 recruitment to CXCR4 with an unchanged potency and basal activity levels (**Fig.1c-d, S1f**). This indicates that physiological cholesterol levels are sufficient for maximum receptor activation, which is not affected by additional membrane cholesterol.

Since cholesterol strongly regulated ligand-induced β-arrestin-2 recruitment to CXCR4, we next investigated its effect on the related atypical receptor ACKR3. Surprisingly, HPβC-mediated reduction of membrane cholesterol resulted in a significantly higher CXCL12-induced β-arrestin-2 recruitment to ACKR3, while restoration of cholesterol levels led to a significant reduction in receptor efficacy (**Fig.1e-g, S1f**). To further determine the influence of cholesterol on receptor activity, we employed CXCR1, CXCR2 and CXCR3. CXCL8-driven efficacy of CXCR1 was partially reduced in lower cholesterol levels, while CXCR2 showed a shift to lower potency, suggesting a compromise in ligand binding (**Fig.1e-g, S1f**). Conversely, CXCR3 showed a similar tendency to that of ACKR3 in its regulation by cholesterol (**Fig.1e-g, S1f**). This contrasts with the role cholesterol plays in the regulation of CXCR4 activity, indicating that cholesterol may act as a positive or a negative allosteric modulator in a receptor-specific manner.

As ligand-induced receptor activation was differentially regulated by cholesterol, we next analyzed its effect on receptor internalization. For this, we utilized a BRET-based system to detect co-localization of Rluc-tagged receptors to the fluorescently tagged membrane marker mem-citrine (**Fig.1h**). Upon stimulation with CXCL12, CXCR4 underwent endocytosis away from the plasma membrane (**Fig.1i-j**). HPβC-mediated decrease in membrane cholesterol levels abolished the ability of CXCR4 to internalize, which was completely restored in the cholesterol-rescued cells. Our results show that CXCL8-induced internalization of CXCR1 and CXCR2 was partially dependent on cholesterol levels, while CXCL12-induced internalization of ACKR3 remained unaffected (**Fig.1i-j**). However, we could not draw conclusions about CXCR3, as CXCL11-induced activation did not trigger significant internalization, likely because of its high degree of constitutive trafficking ^32^.

As many factors are involved in receptor activation, we utilized different controls to examine the regulation of CXCR4 by cholesterol. First, we validated the effect of cholesterol on β-arrestin-2 recruitment in two additional immortalized cell lines, namely HeLa and Cos-7 cells (**Fig.S1g-i**). Next, we evaluated whether cholesterol removal affected basal receptor dimerization levels, as has been described in the literature^18^. For this, we measured BRET between Rluc- and venus-tagged CXCR4 co-expressed in HEK293A cells (**Fig.S1j**). The results show that CXCR4 was able to form homodimers upon cholesterol removal or restoration without significant differences (**Fig.S1k**). Lastly, we examined whether basal subcellular expression levels were affected by the treatments with HPβC. Overall, none of the studied receptors showed significant differences in total expression or subcellular localization between the control and the HPβC-treated groups, with the exception of a higher localization of ACKR3 in the Golgi apparatus (**Fig.S1l-m**). Similarly, restoration of membrane cholesterol levels only caused minor changes in receptor subcellular localization, namely an increase of CXCR4 in the ER, and higher localization of CXCR2 and ACKR3 in the Golgi Apparatus (**Fig.S1l-m**).

### Cholesterol modulates constitutive and ligand-induced G protein signaling

GPCR activation leads to G protein coupling, which triggers canonical downstream signaling readouts, such as proliferation and motility. However, atypical chemokine receptors, such as ACKR3, are only able to recruit β-arrestin. Thus, we continued to evaluate the role of cholesterol in regulating G protein coupling to CXCR1-4.

We first employed a BRET-based mini-G coupling assay consisting of Rluc-tagged receptor and venus-tagged mini-Gα_s/I_ (**Fig.2a**). Our results show that only CXCR4 was significantly affected in its efficacy to couple to mini-Gα_s/i_, while CXCR1, CXCR2 and CXCR3 remained unchanged (**Fig.2b-c, S2a**). In contrast to the complete dependence of CXCR4 on membrane cholesterol to recruit β-arrestin (**Fig.1c-d**), G protein coupling to CXCR4 was only partially dependent on cholesterol levels. While the potency of CXCL11 and CXCL12 was unchanged in the activation of CXCR3 and CXCR4, CXCL8 exhibited a weaker potency to activate CXCR1 and CXCR2 in lower membrane cholesterol levels (**Fig.2b,d, S2a**). This resembles the negative effect on CXCR2 potency observed for β-arrestin recruitment (**Fig.1g**) and suggests that cholesterol may act specifically on these receptors to influence ligand binding.

Upon coupling to active receptors, G proteins undergo activation, leading to heterotrimer dissociation and subsequent intracellular signaling. To understand how cholesterol influences receptor-driven G protein activation, BRET was measured upon binding of fluorescently labeled Gβγ to nanoluciferase (Nluc) fused to the C-terminal tail of GRK3^33^ (**Fig.2e**). The results revealed that constitutive activity (E_min_) levels for CXCR1, CXCR2 and CXCR4, but not CXCR3, decreased upon treatment with HPβC (**Fig.2f-g**), while maximal efficacy (E_max_) remained unchanged in all cases (**Fig.S2b**). This contrasts with the reduction in both maximal efficacy (**Fig.2c**) and constitutive activity (**Fig.S2a**), observed for mini-Gα_s/i_ coupling to CXCR4 with reduced cholesterol levels. Importantly, CXCR1 and CXCR2, but not CXCR3 and CXCR4, showed significantly weaker potencies to respond to CXCL8 with lower membrane cholesterol (**Fig.2h**), similarly to mini-Gα_s/i_ (**Fig.2d**).

The characterization of the effect of cholesterol on β-arrestin recruitment, receptor internalization and G protein signaling shows a large degree of receptor selectivity. While CXCR3 remained unaffected in all readouts, reduced amount of membrane cholesterol resulted in significantly lower activation of CXCR1, CXCR2, CXCR4, or higher ACKR3 activity (**Fig.2i**).

### Interaction with cholesterol dictates constitutive signaling levels in CXCR4

Ligand-induced activation of CXCR4 is strongly compromised by reduced membrane cholesterol levels. To better understand how basal CXCR4 activity influences cholesterol dependence we mutated Asn^3.35^ to a serine (N119S), which has been previously reported to increase the receptor’s constitutive activity^22^. Compared to wild-type (WT) CXCR4, this mutation slightly increased total receptor expression (**Fig.S3a**), while strongly reducing surface levels, possibly due to increased basal receptor internalization (**Fig.S3b**). Both WT and N119S were able to recruit β-arrestin-2 with a similar maximal activation (E_max_), although N119S responded to CXCL12 with a stronger potency (**Fig.3a-c**). Notably, low cholesterol levels compromised the E_max_ of WT CXCR4, but not that of N119S, which retained complete activity in response to CXCL12 with unchanged potency (**Fig.3a-c**). Constitutive recruitment β-arrestin-2 to both WT CXCR4 and N119S remained unaltered in all cases.

We next examined the effect of N119S in mini-Gα_s/i_ coupling. In this case, the effects of cholesterol on N119S constitutive activity (E_min_) levels were similar to those detected for WT CXCR4 (**Fig.3d-e**). As observed for β-arrestin-2, neither the potency nor the ligand-induced E_max_ of N119S were affected by lower membrane cholesterol (**Fig.3f, S3c**). This suggests that ligand-induced activation of N119S does not depend on cholesterol levels.

In high-resolution structures of chemokine receptors, cholesterol has been repeatedly shown to interact directly with Trp^4.50^, which is conserved in 94% of all Class A GPCRs^19,26^. Thus, we mutated this residue to a tyrosine (W161Y) to examine its role in the effect of cholesterol on receptor signaling. This mutant resulted in higher receptor expression, while maintaining a similar surface expression level to that of WT CXCR4 (**Fig.S3a-b**). Measuring β-arrestin-2 recruitment, we observed that low cholesterol levels only reduced W161Y activity by ≈45%, in contrast to a loss of >85% in WT CXCR4 (**Fig.3a-b**). In both cases potency for CXCL12 and baseline recruitment remained unaffected by cholesterol levels (**Fig.3c**). Importantly, W161Y was resistant to low membrane cholesterol for constitutive coupling (E_min_) of mini-Gα_s/i_ (**Fig.3d-e, S3c**). This contrasts with the strong dependence of WT CXCR4 and N119S on cholesterol for constitutive G protein coupling. Finally, we observed that low cholesterol affected ligand-induced (E_max_) mini-Gα_s/i_ coupling to both W161Y and WT CXCR4 (**Fig.3f, S3c**). This suggests that constitutive G protein coupling to CXCR4 may be heavily influenced by specific contacts with cholesterol in the membrane.

Based on these results, we next wanted to determine if cholesterol plays a direct role in receptor activation in the absence of potential effects of the cellular environment. To do so, we examined the effect of cholesterol on CXCR4 activity using an *in vitro* binding assay with purified GFP-tagged mini-Gα_o_ and purified CXCR4 embedded in nanodiscs (**Fig.3g**). Since nanodiscs allow for customization of lipid composition, we assayed CXCR4 in an environment containing 1-palmitoyl-2-oleoyl-*sn*-glycero-3-phosphocholine (POPC) with or without 20% cholesterol. The results showed that WT CXCR4 had a higher coupling to mini-Gα_o_ in the presence of cholesterol (**Fig.3h**). This contrasts with N119S, whose higher constitutive activity masked regulation by cholesterol, and W161Y, whose coupling to mini-Gα_o_ was not increased by cholesterol (**Fig.3h**). This resembles the regulation observed in our cell-based assays (**Fig.3e**). While cholesterol increased CXCL12-induced *in vitro* mini-Gα_o_ binding to WT CXCR4, it failed to do so in either N119S or W161Y (**Fig.3i**). In all cases, we employed IT1t as a control to displace CXCL12-induced activation of CXCR4. In summary, cholesterol regulation of CXCR4 and the mutants thereof in nanodisc systems is consistent with our cell-based readouts, showing that the effects are directly on the receptor and independent of the cellular environment.

### Cholesterol facilitates the transition of agonist-bound CXCR4 to active states

smFRET has been previously utilized to study the dynamic changes between the active and inactive states of CXCR4 in response to CXCL12 and IT1t^30^. We decided to use a similar approach to study how cholesterol influences receptor activation status. To monitor conformational dynamics of CXCR4 using smFRET, we introduced cysteine mutations at residues 150^4.40^ in transmembrane helix 4 (TM4) and 233^6.29^ in TM6 to enable covalent fluorophore labeling using the donor Alexa Fluor 555 (A555) and the acceptor Cyanine5 (Cy5). Purified CXCR4 molecules were then incorporated into POPC nanodiscs, with or without 20% cholesterol, and immobilized on a quartz surface via biotinylation of the C-terminal AviTag™ on CXCR4 (**Fig.4a-b**).

Using apo-CXCR4, we observed three distinct receptor activation states in the absence of cholesterol: 34% low-FRET active state (R*, *E_app_*≈ 0.30), 43% mid-FRET inactive-like intermediate state (R’, *E_app_* ≈ 0.73), and 23% high-FRET inactive state (R, *E_app_* ≈ 0.91) (**Fig.4c, Table S1**). These states closely resemble the ones reported previously^30^. To better understand transitions between FRET states, we next generated two-dimensional transition density probability (TDP) plots. The data showed the existence of bidirectional transitions of CXCR4 molecules between the inactive R and R’ states, while transitions from or to the active R* state were rarely observed (**Fig.4c, Table S1**). Addition of cholesterol allowed the formation of four FRET states (R*, R*’, R’ and R), favoring the inactive R state over the active R*, active-like R*’ and the inactive-like R’ states. (**Fig.4d**).

We next examined the effect of CXCL12 on receptor activation states. In contrast to apo-CXCR4, CXCL12 resulted in a main active R* state (47%), while only 15% of the molecules remained in the inactive R state (**Fig.4e, Table S1**). Addition of CXCL12 also created an active-like state (R*’, *E_app_*≈ 0.52) that replaced the inactive-like R’ state observed with apo-CXCR4. The TDP plots indicated a sequential transition between R and R*’ states, and between R*’ and R states, which rarely occurred directly between R and R* states (**Fig.4e**).

Addition of cholesterol strongly influenced the dynamics of CXCR4 molecules by favoring active receptor conformations (**Fig.4f, Table S1**). As observed with apo-CXCR4, cholesterol allowed the dynamic formation of four distinct FRET states (R*, R*’, R’ and R), which mainly transitioned between the active R* and active-like R*’ states, and rarely between active (R* and R*’) and inactive molecules (R and R’) (**Fig.4f**). The presence of unique conformational states and the differences in transition frequencies between them may explain the effects of membrane cholesterol levels on the efficacy detected in our signaling assays (**Figs.1-3**).

**Fig. 1:**
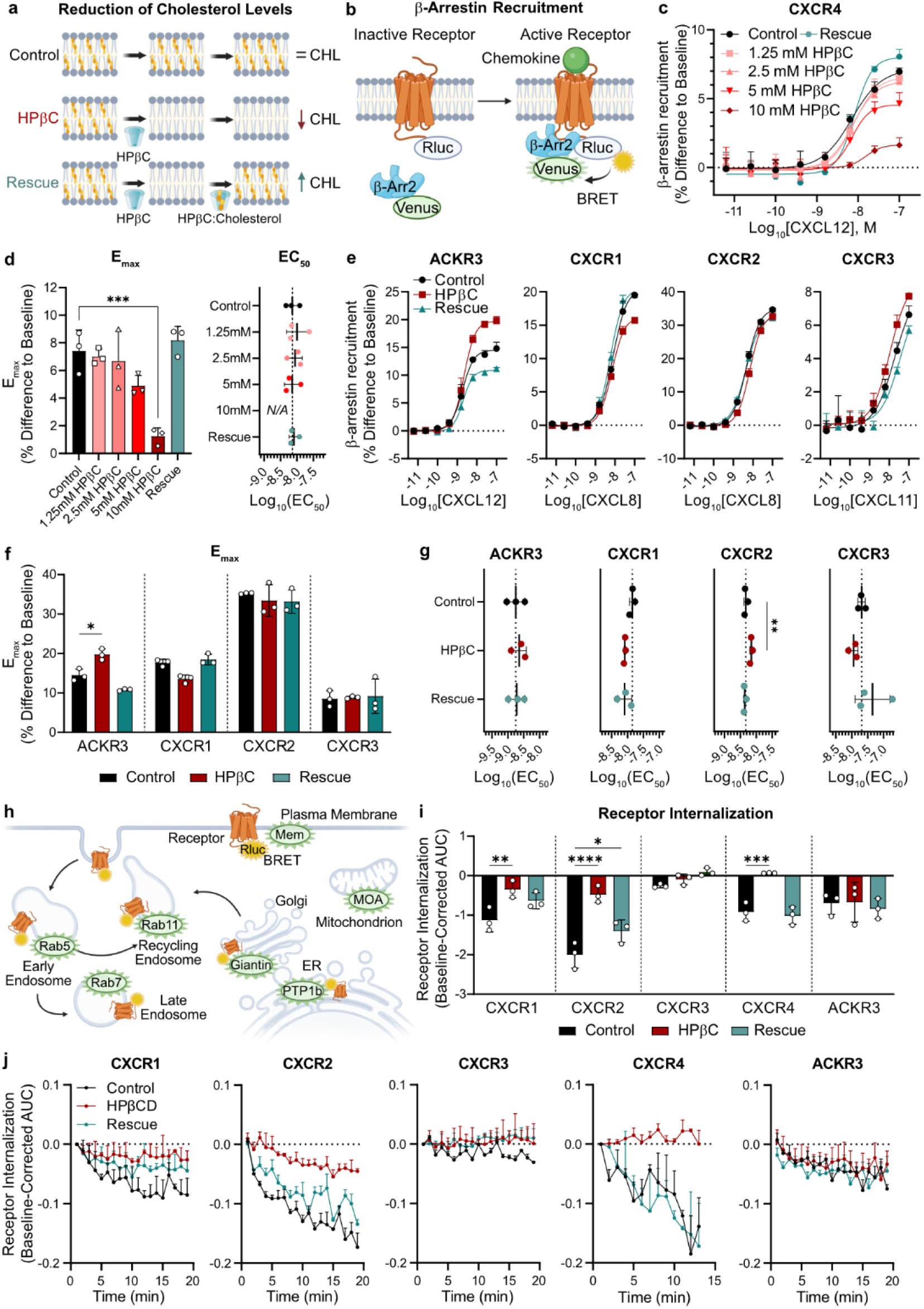
Cholesterol selectively modulates β-arrestin recruitment and internalization of chemokine receptors. **a**, Treatments employed to decrease the amount of membrane cholesterol using hydroxypropyl–β–cyclodextrin (HPβC). In all figures, the rescue group denotes a first treatment with 10mM HPβC, followed by a second treatment with cholesterol–loaded HPβC at a 1mM:10mM ratio (cholesterol:HPβC). **b**, Assay scheme for the direct recruitment of N-terminally Venus–tagged β–arrestin2 to C–terminally Rluc3–tagged CXCR4 using bioluminescence resonance energy transfer (BRET1). **c**, Dose–response curves for CXCL12–induced β–arrestin2 recruitment to CXCR4 in cells treated with the indicated concentrations of HPβC. **d**, Quantification of the effects of decreased cholesterol on the efficacy (E_max_) (left) and the potency (EC_50_) (right) for the recruitment of β–arrestin2 to CXCR4. **e**,**f**,**g**, Effect of decreased cholesterol on the recruitment of β–arrestin2 to CXCR1, CXCR2, CXCR3 and ACKR3. Changes in the dose–response curves (**e**), the efficacy (**f**), and the potency (**g**) for the recruitment of β–arrestin2 to each receptor. **h**, Scheme for the direct detection of the co–localization of CXCR1, CXCR2, CXCR3, CXCR4 and ACKR3 with the indicated Venus–tagged subcellular localization markers. **i**,**j**, Changes in ligand-induced receptor internalization upon reduction of cholesterol levels, shown as area under the curve (**i**), and time–resolved BRET signals (**j**). CXCR1 and CXCR2 were stimulated with 50 nM CXCL8, CXCR3 was stimulated with 50 nM CXCL11, and CXCR4 and ACKR3 were stimulated with 50 nM CXCL12. (**b**,**h**) Created with BioRender.com. (**c-g**,**i-j**) The asterisk symbols indicate statistically significant differences (*p < 0.05, **p < 0.01, ***p < 0.001, ****p < 0.0001) between the indicated groups, determined by one-way ANOVA. N = 3 independent experiments, performed in technical triplicates. All data are shown as mean ± SD. Only significant differences to the control group are shown.

**Fig. 2:**
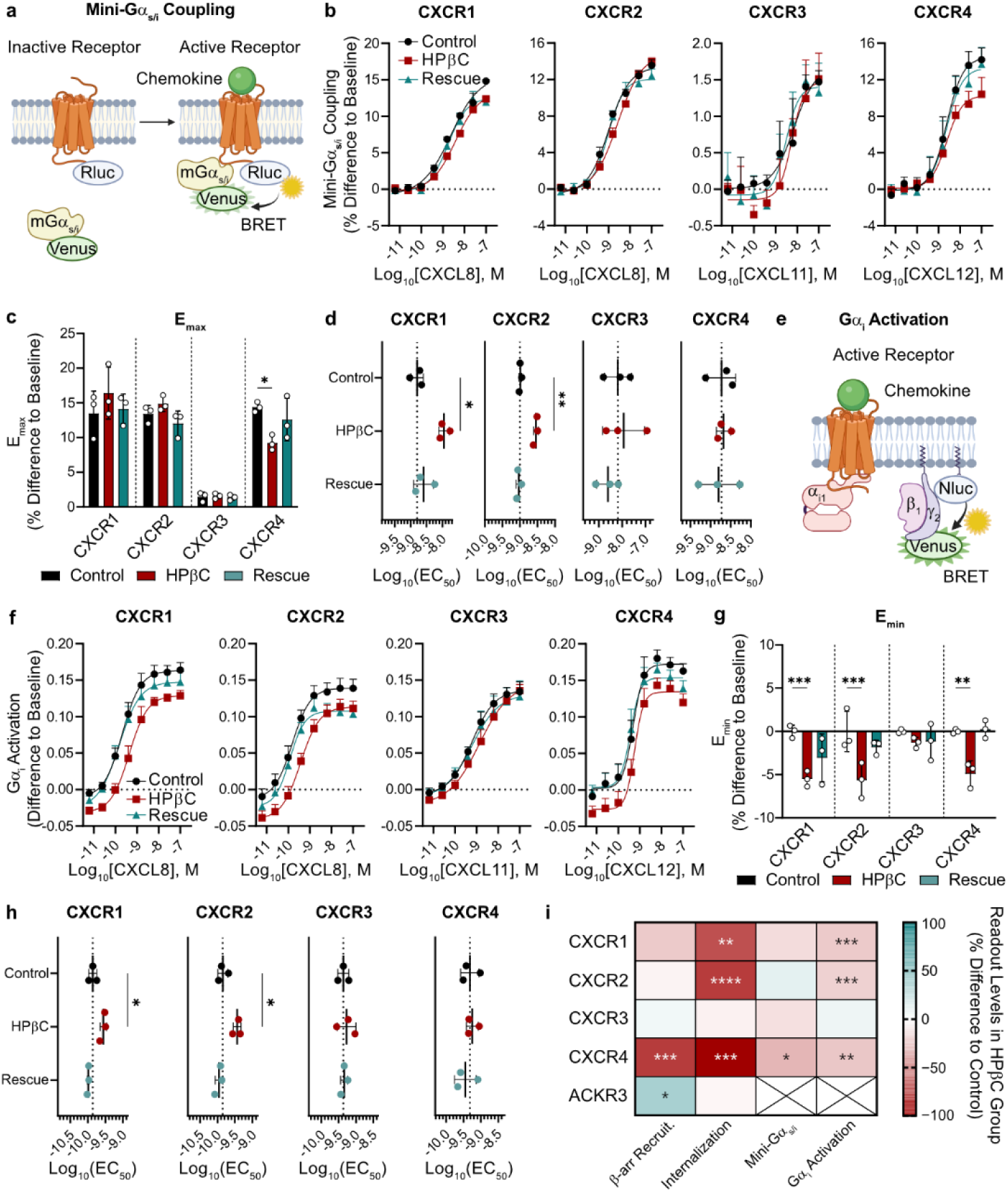
Cholesterol modulates constitutive and ligand-induced G protein signaling in chemokine receptors. **a**, Schematic representation of the BRET-based direct mini-Gα_s/i_ coupling assay used to study CXCR1, CXCR2, CXCR3 and CXCR4. **b**,**c**,**d**, Effect of decreased cholesterol using HPβC on ligand-induced mini-Gα_s/i_ coupling. The results are shown as dose-response curves (**b**), and the quantification of the efficacy (**c**), and the potency (**d**) for mini-Gα_s/i_ coupling to each receptor. **e**, Scheme of the heterotrimer dissociation assay (from now on referred to as Gα_i_ activation) employed to determine the role of decreased cholesterol in constitutive G protein signaling enacted by CXCR1, CXCR2, CXCR3 and CXCR4. BRET signal was detected upon the translocation of the β_1_ and Venus-tagged γ_2_ subunits G protein subunits to the plasma membrane, where they came into proximity to the NanoLuc® luciferase (Nluc)-tagged C-terminal tail of GRK3. Cells were also transfected with untagged α_i1_ subunit and untagged receptor. **f**,**g**,**h**, Changes in constitutive Gα_i_ activation levels upon reduction of the amount of membrane cholesterol using HPβC, shown as dose-response curves (**f**), as well as efficacy (**g**) and potency differences (**h**). **i**, Heat-map summarizing the effects of HPβC–mediated cholesterol extraction on the efficacy of the four studied readouts triggered by chemokine receptor activation. (**a**,**e**) Created with BioRender.com. (**b-d**,**f-i**) The asterisk symbols indicate statistically significant differences (*p < 0.05, **p < 0.01, ***p < 0.001, ****p < 0.0001) between the indicated groups, determined by one-way ANOVA. N = 3 independent experiments, performed in technical triplicates. All data are shown as mean ± SD. Only significant differences to the control group are shown.

**Fig. 3:**
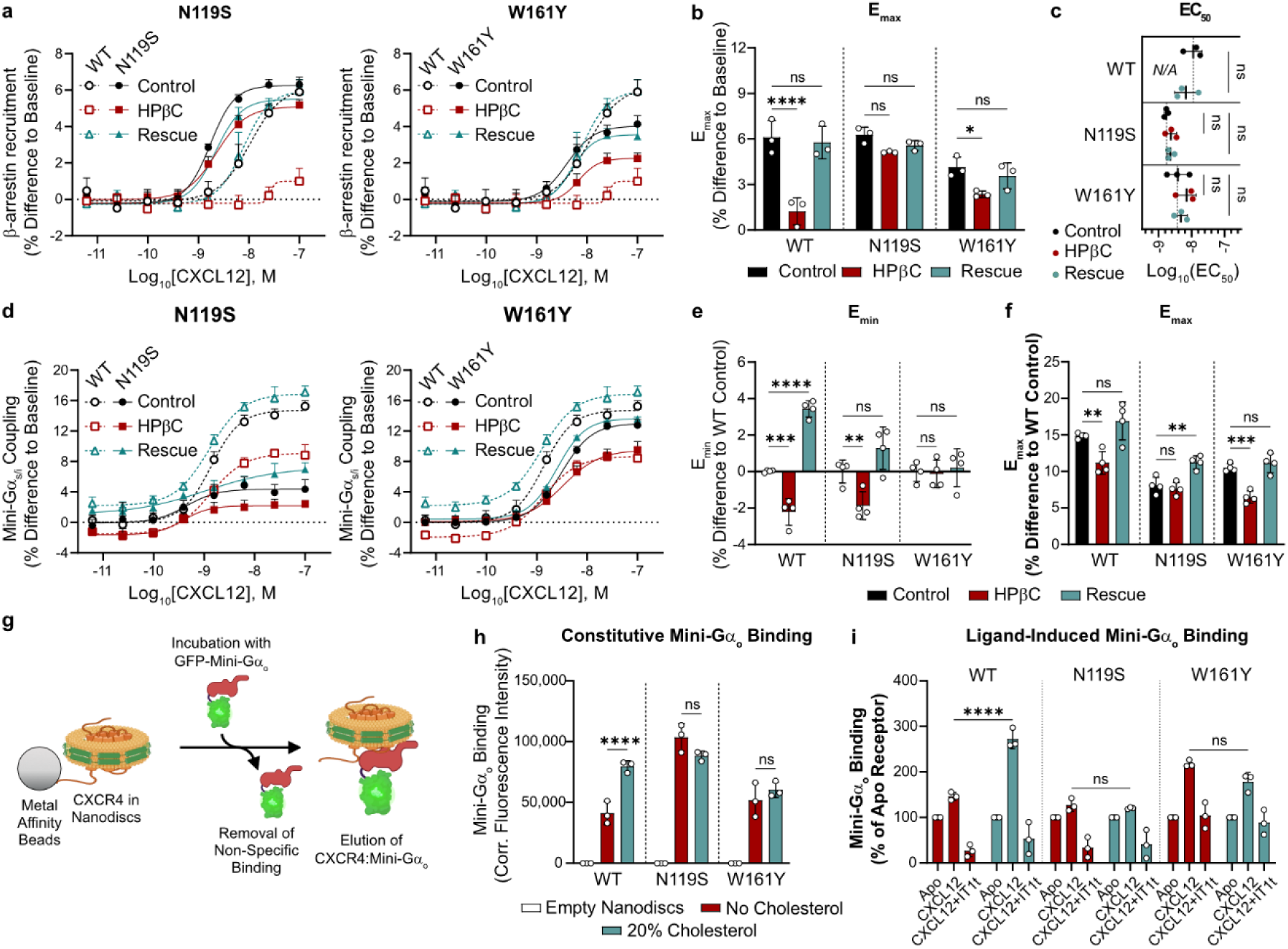
Cholesterol facilitates the activation of CXCR4, enabling efficient G protein coupling. **a**,**b**,**c**, Effect of decreased cholesterol on the recruitment of N-terminally Venus-tagged β–arrestin2 to the C-terminally Rluc3-tagged CXCR4 mutants N119S and W161Y, compared to wildtype (WT) CXCR4. The results are shown as dose-response curves (**a**), efficacy (**b**), and potency (**c**). **d**,**e**,**f**, Changes in mini-Gα_s/i_ coupling to WT CXCR4 and its mutants in cholesterol-decreased or rescued HEK293A cells. The dose-response BRET signal of C-terminally Rluc3-tagged WT CXCR4 was compared to that of N119S (**d**, left) and W161Y (**d**, right), which was used to determine the constitutive (**e**) and maximal ligand-induced (**f**) coupling to N-terminally Venus-tagged mini-Gα_s/i_. **g**, Schematic representation of the *in vitro* fluorescent saturation binding assay (FSBA) performed with C-terminally His-tagged CXCR4 MSP1D1-nanodics and N-terminally GFP-tagged mini-Gα_o_. Created with BioRender.com. **h**,**i**, Quantification of the constitutive (**h**) and ligand-induced (**i**) levels of mini-Gα_o_ coupling to WT CXCR4, N119S and W161Y. CXCL12 was displaced by the inverse agonist IT1t as a control for lack of mini-Gα_o_ binding to CXCR4. (**a-f,h-i**) The asterisk symbols indicate statistically significant differences (*p < 0.05, **p < 0.01, ***p < 0.001, ****p < 0.0001, ns denotes non-significant changes) between the indicated groups, determined by one-way ANOVA. N = 3 independent experiments, performed in technical triplicates. All data are shown as mean ± SD.

**Fig. 4:**
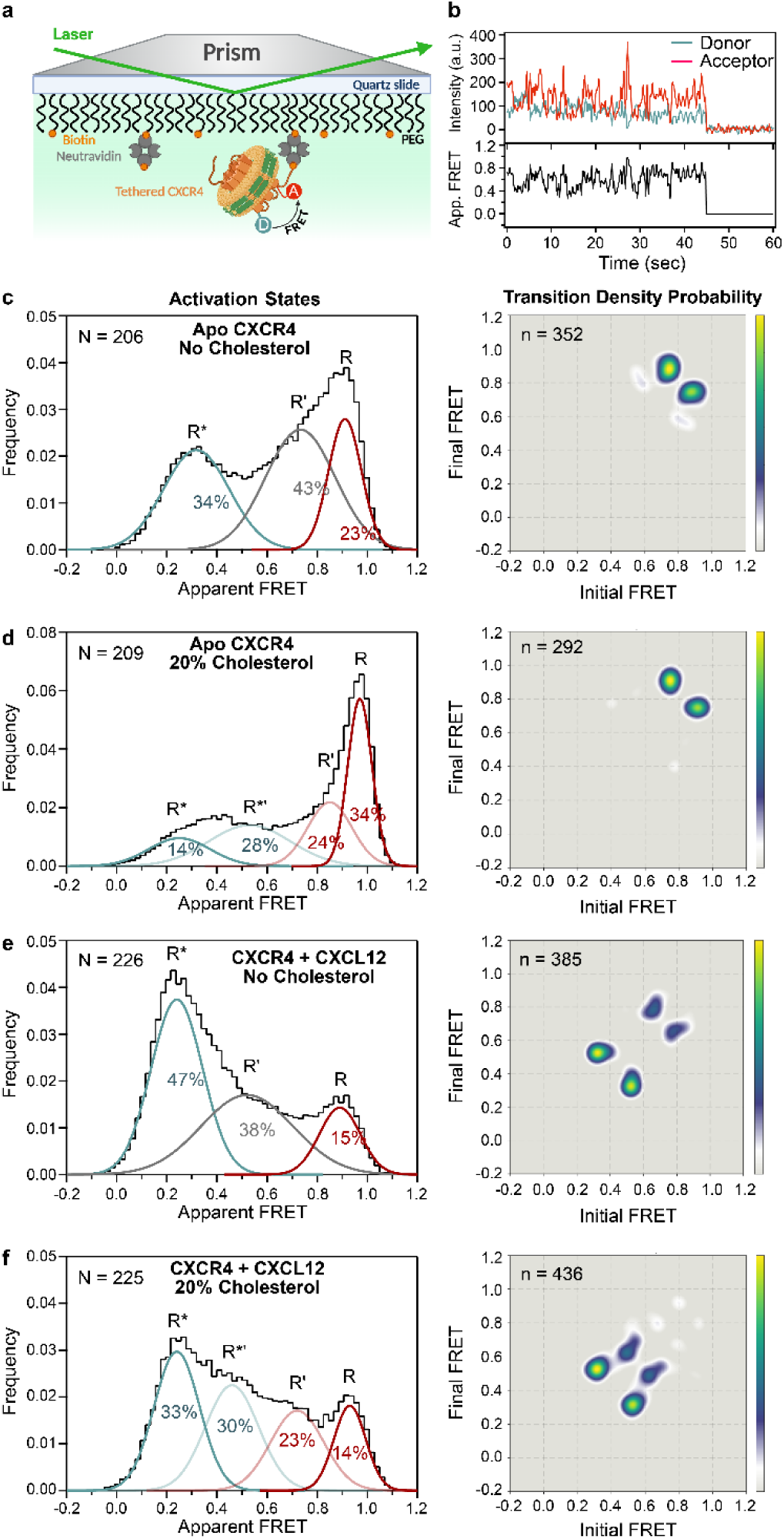
Cholesterol facilitates ligand-induced transitions of CXCR4 to active states. **a**, An individual receptor (orange) was tagged with donor and acceptor fluorescent dyes, incorporated into a phospholipid (yellow) nanodisc (green), and immobilized to a quartz slide through a biotin (orange circle)–neutravidin (dark gray) linkage. A prism generates total internal reflection of the excitation laser, restricting excitation to donor fluorophores located near the surface. Created with BioRender.com. **b**, Representative single-molecule time traces for apo-CXCR4; donor (blue) and acceptor (red) intensities are shown in the top panel, and the corresponding apparent FRET efficiency (black) is shown in the bottom panel. **c-f**, Changes in the activation states of ligand-free CXCR4 in the absence (**c**) and presence (**d**) of 20% cholesterol; or CXCL12-bound CXCR4 in the absence (**e**) or presence (**f**) of 20% cholesterol using single–molecule fluorescence resonance energy transfer (smFRET). Left panels show histograms of single-molecule FRET states, while the right panels show two-dimensional transition density probability (TDP) plots for the transitions between observed FRET states. N represents the number of single-molecule traces used to generate histograms, and n represents the total number of transitions across the states.

Finally, we investigated the effect of the inverse agonist IT1t on CXCR4 conformational dynamics. In the absence of cholesterol, receptor activation states and the transitions between them were similar to those detected in apo-CXCR4 (**Fig.S4a, Table S1**). Addition of cholesterol to IT1t-bound CXCR4 created an active-like R*’ state, as previously detected for apo and CXCL12-bound receptors and shifted the receptor to more active conformations (**Fig.S4b, Table S1**), resembling the effect of cholesterol on CXCL12-bound CXCR4 (**Fig.4f**). The TDP plots showed that, while the receptor preferred inactive conformations, transitions between the inactive R and inactive-like R’ states occurred more frequently (**Fig.S4b**).

### Oxysterols fail to rescue the effects of cholesterol on chemokine receptor activity

Oxysterols have been recently reported to increase in cellular senescence, where they alter GPCR signaling^34^; however, the mechanisms underlying this dysregulation remain unclear. To examine this further, we utilized two of the most abundant senescence-associated hydroxycholesterol (HCHL) species: 25-HCHL and 7β-HCHL (**Fig.5a**). Upon HPβC-mediated reduction in cholesterol levels, HCHLs were added to the cells. While CXCL12 induced β-arrestin-2 recruitment to CXCR4 expressed in control and cholesterol-rescued cells, both 25-HCHL and 7β-HCHL failed to restore receptor activity (**Fig.5b**).

As 25-HCHL appeared to completely abolish β-arrestin-2 recruitment to CXCR4, we investigated its effects on membrane composition. While the addition of 25-HCHL significantly increased cellular sterol levels, it surprisingly reduced membrane rigidity without affecting cell viability (**Fig.S5a-c**). Among the other CXC-type chemokine receptors, 25-HCHL only reduced efficacy in CXCR1; however, it further weakened the potency of CXCR2 activation by CXCL8 (**Fig.5c, Fig.S5d**). Surprisingly, 25-HCHL selectively strengthened the potency of CXCL11 for CXCR3 (**Fig.5c**).

We next examined the effect of 25-HCHL in G protein coupling and activation. The results showed that the efficacy of ligand-induced mini-Gα_s/i_ coupling was not significantly affected, while 25-HCHL addition resulted in weaker potency for CXCL8 to activate both CXCR1 and CXCR2, compared to the control cells (**Fig.5d, Fig.S5e**). Finally, we evaluated how 25-HCHL influenced G protein activation. This oxysterol failed to restore constitutive activity in any of the studied receptors and reduced the potency of CXCR1 and CXCR2 to respond to CXCL8 (**Fig. 5e**, **Fig. S5f-g**).

**Fig. 5:**
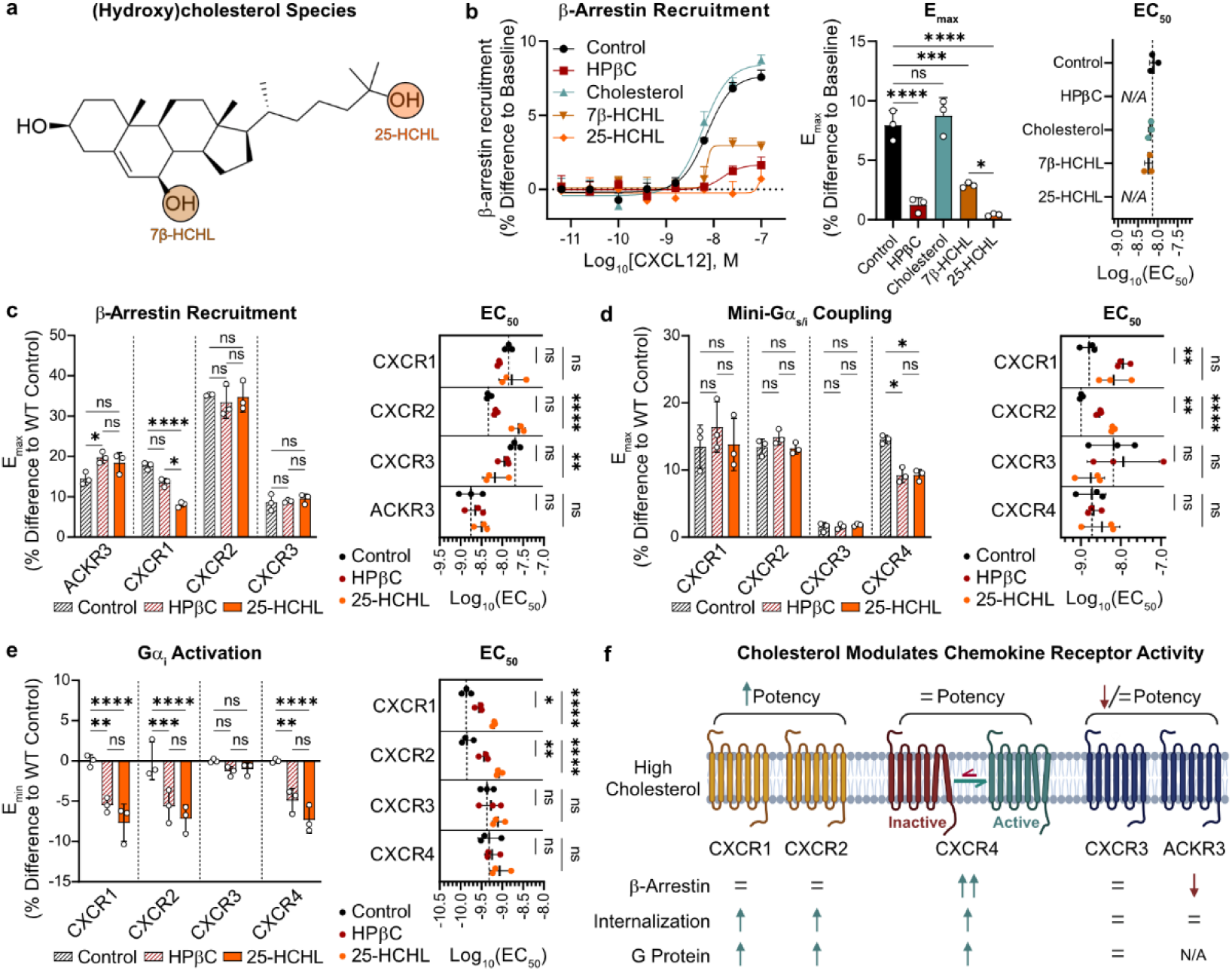
Oxysterols fail to rescue the effects of cholesterol on chemokine receptor activity. **a**, Molecular structure of cholesterol, showing the hydroxylation sites in 25-hydroxycholesterol (HCHL) and 7β-HCHL. **b**, Changes in N-terminally Venus-tagged β–arrestin2 recruitment to C-terminally Rluc3-tagged CXCR4 in HEK293A cells rescued with HPβC preloaded with 25-HCHL and 7β-HCHL. Changes in dose–response recruitment (left), efficacy (middle), and potency (right). **c**, Effect of 25-HCHL addition on the recruitment of β–arrestin2 to chemokine receptors, for which efficacy (left), and potency (right) were calculated. **d**, Changes in the efficacy (left), and the potency (right) for N-terminally Venus-tagged mini-Gα_s/i_ coupling to C-terminally Rluc3-tagged chemokine receptors rescued with 25-HCHL. **e**, Role of 25-HCHL in the constitutive Gα_i_ activation levels of untagged chemokine receptors, calculated as changes in efficacy (left), and potency (right). **f**, Summarizing scheme of the role of decreased cholesterol and oxysterol addition in chemokine receptor ligand binding and intracellular signal transduction; created with BioRender.com. (**b-e**) The asterisk symbols indicate statistically significant differences (*p < 0.05, **p < 0.01, ***p < 0.001, ****p < 0.0001, ns denotes non-significant changes) between the indicated groups, determined by one-way ANOVA. N = 3 independent experiments, performed in technical triplicates. All data are shown as mean ± SD.

Overall, the signaling results for CXCR1-4 and ACKR3 can be classified as three distinct modes of cholesterol-mediated regulation of receptor activity (**Fig.5f**). CXCR1 and CXCR2 were only partially affected by cholesterol reduction and showed weaker potency for CXCL8, while CXCR4 displayed a strong dependence on cholesterol for β-arrestin recruitment, receptor internalization, and G protein signaling. In contrast, ACKR3 and CXCR3 exhibited enhanced activity and potency under reduced cholesterol levels.

## Discussion

Membrane lipid composition strongly regulates GPCR function, and many high-resolution structures of GPCRs contain sterols and other lipids bound to the receptor TM domain. However, experiments to assess the specific roles of individual lipid species at a molecular level and the functional importance of receptor-associated sterols have been technically challenging, limiting our mechanistic understanding of cholesterol regulation. Here, we isolated the role of cholesterol using HPβC-mediated extraction in mammalian cells and correlated the effects on receptor function with modulation of receptor conformation and G protein binding in nanodiscs. By comparing multiple chemokine receptors and cholesterol derivatives, we show that this regulation is receptor-, pathway- and sterol-specific and driven by stabilization of specific conformational states, revealing mechanistic insights into the regulation of GPCRs by cholesterol.

Comparison of five CXC-type chemokine receptors revealed that cholesterol regulates receptor activity in a receptor-specific manner, causing three distinct types of pharmacological effects (**Fig.5f**). These findings are consistent with the distinct modes of regulation observed for cholesterol across class A GPCRs^35^. Similar to CXCR4, cholesterol depletion has been shown to reduce signaling in receptors such as the 5-HT1A receptor^36^, A_2A_ adenosine receptor^3^, and μ-opioid receptor, whereas the δ-opioid receptor is comparatively insensitive^37^. In contrast, the β2-adrenergic receptor exhibits reduced signaling upon cholesterol enrichment^38^, resembling our findings for ACKR3. Understanding how cholesterol causes receptor-specific alterations as a positive or negative modulator could have important implications for disease pathophysiology and apply to many class A GPCRs.

Our experiments using CXCR4 mutants provide further insights into the mechanistic role of cholesterol in receptor regulation. While low membrane cholesterol impaired the ability of WT CXCR4 to recruit β-arrestin and couple G proteins, W^4.50^161Y fully or partially bypassed this dependence for constitutive and ligand-induced activity, respectively. This supports a role for Trp^4.50^ as a hub for cholesterol regulation. Given the widespread presence of sterols associated with Trp^4.50^ in GPCR structures, cholesterol binding to this residue may have a similar role in regulating the activity of other class A GPCRs.

Receptors with high constitutive activity were functionally independent of cholesterol-mediated regulation. We show that ligand-induced activation of N^3.35^119S was insensitive to cholesterol. As mutation of Asn^3.35^ disrupts the conserved sodium binding site present in inactive conformations, this suggests that the effects of cholesterol are redundant for a receptor already favoring active conformational states. Interestingly, we also detected that the highly constitutively active receptors CXCR3^32^ and ACKR3^39^ were both functionally independent of cholesterol. Compared to CXCR4, ACKR3 has a higher population of active conformations in the absence of agonist, which drives its high basal activity and may contribute to its lack of sensitivity to cholesterol^30^. Taken together, this suggests that the receptor-specific effects of cholesterol on chemokine receptors are linked to their conformational equilibria and the propensity to populate active conformational states.

Our smFRET measurements provided a molecular mechanism underlying cholesterol-driven modulation of the conformational states of CXCR4. In the absence of cholesterol, CXCL12-bound receptors exist in both inactive and intermediate states with limited transitions to fully active conformations. Cholesterol shifted the equilibrium toward active states, reduced back-transitions to inactive conformations and stabilized a unique, active-like, receptor state. In contrast, cholesterol had more modest effects on apo or inverse agonist-bound receptors, where increased cholesterol favored inactive and active states over intermediate conformations. Collectively, our smFRET results explain why we observed unchanged potency but increased CXCR4 efficacy: productive signaling states are more populated and longer-lived. The stabilization of active or active-like receptor conformations is consistent with the effects on conformational equilibria of the A_2A_ adenosine- and ghrelin receptors,^3,40^ detected by ^19^F-NMR and fluorescence spectroscopy, respectively. Together, these studies point to cholesterol as a key determinant of GPCR conformational dynamics through stabilization of specific conformational states.

Our results with CXCR4 mutants in mammalian cells can be recapitulated with receptors reconstituted into nanodiscs, showing that the effects of cholesterol are directly on the receptors and independent of the cellular environment affecting receptor organization and/or coupling to effector proteins. In addition, the W161Y mutant suggests that specific cholesterol binding sites contribute to the regulation. However, we cannot conclusively uncouple the contributions of specific cholesterol binding from effects on membrane order, which could contribute to alterations of the receptor conformational equilibrium in both cells and nanodiscs. Future studies using defined lipid compositions, including high-resolution structures in the absence and presence of cholesterol will be required to distinguish indirect membrane effects from direct cholesterol-receptor interactions and provide a complete mechanistic understanding of the molecular details underlying cholesterol regulation.

The treatment with 10 mM HPβC reduced rather than eliminated membrane cholesterol, allowing receptor regulation to be studied within a physiologically relevant range. Because cholesterol abundance varies across tissues and intracellular membranes^41^, our results suggest that chemokine receptor signaling may differ depending on the cellular context. For example, CXCR1 and CXCR2, which are highly expressed in neutrophils^9^, would be sensitive to changes in cholesterol availability. Thus, hypocholesterolemia would potentially impair neutrophil chemotaxis and antimicrobial responses, while hypercholesterolemia may enhance inflammatory neutrophil recruitment via increased CXCR1-2 activity. In contrast, the signaling of CXCR3, upregulated in activated CD8⁺ T cells^42^, would potentially remain unaffected by changing cholesterol levels. This suggests that altered cholesterol metabolism may selectively shift the balance between innate and adaptive immunity, which is known to change susceptibility to infection and chronic inflammation^43,44^.

Despite partially restoring membrane sterol content, oxysterols failed to rescue the function of cholesterol in CXC-type chemokine receptors, acting as neutral or negative modulators of receptor activity. This indicates that not all sterol-like molecules share the same functional mode of binding required for GPCR modulation. Cholesterol hydroxylation may disrupt the Van der Waals interactions involved in cholesterol dimerization within the lipid bilayer^45^, thereby causing our observed reduction in membrane rigidity with 25-HCHL. Importantly, these findings link oxidative lipid species produced in senescence or inflammation to dysfunctional signaling in selective receptors, as previously proposed for CXCR2^46^, and CXCR4^34^. Given that CXCR4 frequently dominates chemokine-guided migration, oxysterol-associated reductions in CXCR4 activity may shift immune responses to alternative chemokine gradients during chronic inflammation and aging.

Overall, our work shows that cholesterol not only modifies membrane organization but also acts as a receptor-specific allosteric regulator, with strong implications for lipid oxidation in inflammatory signaling.

## Methods

### Cell culture

Parental HEK293A cells (Thermo Fisher Scientific, Cat#R70507), Chinese hamster ovary (CHO) cells, and COS-7 cells were cultured at 37 °C in a humidified incubator with 5% CO₂ using high-glucose Dulbecco’s Modified Eagle Medium (DMEM) containing GlutaMAX (Thermo Fisher Scientific, Cat#31966), supplemented with 10% fetal bovine serum (FBS) and 1% penicillin–streptomycin (Merck, Cat#P4333). Cells were routinely passaged by washing with phosphate-buffered saline (PBS) followed by detachment using 0.05% trypsin–EDTA (VWR, Cat#392-O459P). *Spodoptera frugiperda* (Sf9) insect cells (Expression Systems, Cat#94-001F) were cultured in suspension in ESF 921 Protein-Free Insect Cell Culture Medium (Clinisciences, Cat#96-001-01) with shaking at 140 rpm, and virus production was performed as previously described^28^.

### Cloning and receptor mutagenesis

The cDNA encoding human C-terminally Rluc3-tagged CXCR1, CXCR2, CXCR3, CXCR4, ACKR3, and the mutants of CXCR4 were expressed in the pcDNA3.1 vector. N-terminally FLAG-SNAP-tagged CXCR1-3^32^, CXCR4^39^, and ACKR3^39^, and the mutants thereof were expressed in the pcDNA5.1 vector. CXCR4 mutants were generated by mutation to the desired amino acid using the Stratagene QuikChange protocol using Phusion High-Fidelity PCR Master Mix with HF Buffer (Thermo Fisher, Cat#F531S) and Dpnl (Bionordika, Cat#R0176S). CXCR1-Rluc3, CXCR2-Rluc3, CXCR3-Rluc3, CXCR4-Rluc8, venus-β-arrestin-2, Nluc-CAAX, and GFP-mini-Gα_o_ were generated using the FastCloning method as described in^47^.

Rluc3-tagged CXCR4 and ACKR3 were a kind gift from Nikolaus Heveker (Université de Montréal, QC, Canada) Mem-citrine was obtained from Jonathan Javitch (Columbia University, NY, USA); venus-tagged PTP1b, giantin, Rab5, Rab7, Rab11, MOA,); venus-mini-Gα_s/i_^48^, and the constructs for the heterotrimer dissociation assays (untagged Gα_i1_, β_1_, venus-γ2, and Nluc-tagged C-terminal tail of GRK3)^33^ were provided by Nevin Lambert (Augusta University, GA, USA; and the original untagged mini-Gα_o_ was kindly provided by Arun Shukla. The sequence of all constructs was confirmed by bidirectional sequencing.

### Cholesterol extraction, determination of cholesterol levels, and cell viability

Cholesterol was extracted using (2-Hydroxypropyl)-β-cyclodextrin (HPβC; Sigma-Aldrich, Cat#H107). A stock 100 mM HPβC solution was dissolved in PBS, allowed to homogenize by gentle shaking at RT for up to 2 hrs, and kept at −20 °C for long-term storage. To rescue membrane cholesterol levels, 100 mM HPβC was preloaded with 10 mM 3β-Hydroxy-5-cholestene (cholesterol; Merck, Cat#C3045), 10mM 25-Hydroxycholesterol (25-HCHL; Merck, Cat#H1015), or 10mM 7β-Hydroxycholesterol (7B-HCHL; MedChemExpress, Cat#HY-113341) at an HPβC:cholesterol ratio of 10:1. Cholesterol was allowed to bind to HPβC by overnight incubation at 4 °C in a rotating disk. Excess cholesterol from the resulting mixtures was removed using a 0.45-µm filter unit, and the stock was kept at −20 °C for long-term storage.

In all experiments indicated in the section “BRET-based signaling assays”, transfection media was removed from the wells two hours before assay reading and replaced with BRET buffer (PBS supplemented with 5mM glucose) for the control group, or HPβC diluted in BRET buffer at the indicated concentrations (10 mM, unless specified) for the HPβC-treated groups. Cells were incubated for 1 h at 37 °C with 5% CO₂. The solutions were then removed from the wells, and BRET buffer (for the control and HPβC groups), or cholesterol-loaded HPβC was added. For this, the stock solutions of cholesterol-loaded HPβC were diluted 20-fold in BRET buffer for a final HPβC concentration of 5 mM to prevent additional cholesterol removal. The cells were then incubated for 1 h at 37 °C with 5% CO₂. Upon incubation, the solutions were removed from the wells, which were then washed twice with PBS, after which BRET buffer was added to continue as specified for each assay.

Using the same experimental setup, cell supernatants were collected to determine the amount of extracted cholesterol using the Amplex™ Red Cholesterol Assay Kit (Thermo Fisher, Cat#A12216) following the manufacturer’s indications. In parallel, we tested changes in cell viability upon HPβC-mediated cholesterol extraction or restoration using CellTiter-Glo® Luminescent Cell Viability Assay (Promega, Cat#G7570).

### Determination of membrane rigidity

HEK293A were trypsinized, centrifuged and resuspended at 2 million cells/ml in PBS containing 5mM glucose. Cells were divided into five tubes, each containing 1 ml, and they were treated with 10 mM HPβC (or PBS for the control cells) at 37 °C for 1 h. Cholesterol was then restored in three of the samples by adding PBS (in the control, or the low-cholesterol group), 5 mM HPβC-loaded cholesterol, 25-HCHL, or 7β-HCHL for an additional hour. Upon incubation, excess cholesterol was removed by two centrifugation steps at 200g followed by PBS washes.

Cell membrane rigidity was then measured using the fluorescence anisotropy of the hydrophobic probe 1,6-Diphenyl-1,3,5-hexatriene (DPH, Sigma-Aldrich, Cat#D208000) following an adjusted procedure from Nagy et al.^49^ HEK293A cells were adjusted to 1.2 million cells/mL, washed with PBS, resuspended at 12 million cells/mL in labelling buffer (PBS with 5mM glucose and 2 µM DPH) and agitated at room temperature for 20 minutes. Cells were then diluted to 1.2 million cells/mL with PBS supplemented with 5mM glucose and 200 µL were transferred to a black 96-well plate and immediately placed in spectrophotometer (prewarmed to 37 °C) for measurement. We next measured DPH anisotropy using an excitation of 350 nm and emission of 452 nm. Three measurements were recorded at 0, 5, 10 minutes, where the measurement at 5 minutes was used to determine rigidity, expressed as a raw fluorescence anisotropy ratio.

### Purification of CXCL12, MSP1D1, BirA and GFP-mini-Gα_o_ from bacteria

Human CXCL12, MSP1D1, BirA and GFP-mini-Gα_o_ were expressed and purified following previously published protocols with minor modifications^29,50–52^. Briefly, each protein was expressed in *Escherichia coli* (BL21-Gold (DE3)pLysS) from a pET-21 (for CXCL12), pET28a (for MSP1D1 and BirA), or pET-15b (for GFP-mini-Gα_o_) vector encoding an N-terminal His₈-(for CXCL12 and MSP1D1) or His_6_-tag (for BirA and GFP-mini-Gα_o_) followed by an enterokinase cleavage site (for CXCL12) or a Tobacco Etch Virus (TEV) protease cleavage site (for MSP1D1, BirA and GFP-mini-Gα_o_). After expression, CXCL12 was isolated from inclusion bodies in unfolding conditions, while MSP1D1, BirA and GFP-mini-Gα_o_ were isolated from the cell lysate. Each protein was then purified using Ni-NTA affinity chromatography. For CXCL12, MSP1D1 and GFP-mini-Gα_o_, the His-tag was removed by enterokinase digestion, and the resulting untagged proteins were further purified by a second Ni-NTA affinity step followed by reverse-phase chromatography. Finally, the proteins were buffer-exchanged into 20 mM acetate pH 4.0 (for CXCL12); 20 mM HEPES pH 7.5, 150 mM NaCl and 5 mM sodium cholate (for MSP1D1); 20 mM Tris pH 7.5, 100 mM NaCl (for BirA); or 20mM HEPES pH 7.4, 500mM NaCl, 10% Glycerol, 50μM GDP and 1mM MgCl_2_ (for GFP-mini-Gα_o_). The proteins were then concentrated by centrifugal filtration and stored at −80 °C until use.

### BRET-based signaling assays

For these assays, HEK293A, HeLa, or Cos-7 cells (30,000 cells per well) were seeded in poly-D-lysine-coated white 96-well plates (Greiner. Cat#655083) and simultaneously co-transfected as indicated below for each experiment using PEI at a DNA:PEI ratio of 1:2 (w/v). After 48 hours, the cells were treated as indicated in the section “Cholesterol extraction using cyclodextrin”. Having the cells in BRET buffer, (PBS supplemented with 5mM glucose), the cells were incubated with BRET substrate (Coelenterazine H, Nanolight technology, Cat#301–10) for 8 minutes at a final concentration of 5 μM, after which the cells were stimulated with the ligand, and the plate was then read at 37 °C using an EnVision Plate Reader (Revvity, Inc.) to measure BRET ratios. The ligands for these assays were: CXCL8 (PeproTech®, Cat#200-08M) for CXCR1 and CXCR2, CXCL11 (PeproTech®, Cat#300-46) for CXCR3, and CXCL12 for CXCR4 and ACKR3. BRET values were calculated as follows:

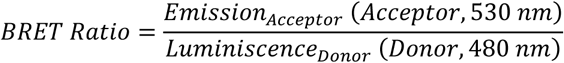

All dose-response curves were fitted with a 4-parameter nonlinear regression model (variable slope) in Graphpad prism (v10.2.3) and efficacy and baseline values were extracted from top and bottom fit, respectively. Potencies were extracted from Log(EC_50_) values of the fits.

The transfected amounts for each type of signaling assay are indicated below:

I. Direct β-arrestin recruitment assay^39^. HEK293A cells were co-transfected with 2.5 ng/well of C-terminally Rluc3-tagged receptor and 10 ng/well venus-β-Arrestin2, using empty pcDNA3.1 vector to a total transfected DNA amount of 50 ng/well. To increase signal intensity, HeLa and Cos-7 cells were transfected with 10 ng/well receptor-Rluc3 and 40 ng/well venus-β-Arrestin2, for a total transfection of 50 ng/well.
II. Receptor homodimerization^53^. HEK293A cells were co-transfected with 0.75 ng/well C-terminally Rluc8-tagged CXCR4 or the membrane marker Nluc-CAAX (used as a negative control), and a concentration range (from 0 to 16 ng/well) of C-terminally venus-tagged CXCR4, where empty pcDNA3.1 was added to a total DNA amount of 50 ng/well.
III. Cell-based mini-Gα_s/i_ coupling^48^. In this case, HEK293A cells were co-transfected with 2.5 ng/well C-terminally Rluc3-tagged CXCR4 and 10 ng/well venus-mini-Gα_s/i_, adding empty pcDNA3.1 for a final DNA amount of 50 ng/well.
IV. Heterotrimer dissociation assay for G protein activation^33^. HEK293A cells were co-transfected with 5 ng/well N-terminally SNAP-tagged receptor, 36 ng/well untagged Gα_i1_, 12.5 ng/well β_1_, 12.5 ng/well venus-γ2, and 2.5 ng/well Nluc-tagged C-terminal tail of GRK3, for a final amount of DNA of 68.5 ng/well.
V. Subcellular localization assay^54^. HEK293A cells were co-transfected with 2.5 ng/well of C-terminally Rluc8-tagged receptor plasmid, 30 ng/well of each venus-tagged subcellular compartment marker, and 17.5 ng of pcDNA3.1 to obtain a total DNA amount of 50 ng/well. To detect total receptor expression levels, 30 ng/well of empty pcDNA3.1 vector was added instead of any labeled subcellular marker. Receptor localization was assessed by measuring the BRET signal between Rluc8-tagged receptors and Venus-tagged organelle markers, enabling the determination of the subcellular distribution of CXCR1, CXCR2, CXCR3, CXCR4, and ACKR3. The compartment-specific markers were mem-citrine^55^ for the plasma membrane, venus-PTP1b^54^ for the endoplasmic reticulum, venus-giantin^54^ for the Golgi apparatus, venus-Rab5a^56^ for early endosomes, venus-Rab7a^57^ for late endosomes, venus-Rab11a^57^ for recycling endosomes, and venus-MOA^54^ for mitochondria.

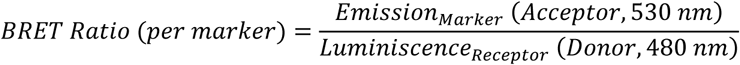

The mitochondrial BRET signal served as a background reference for calculating fold enrichment of receptor localization within each compartment.

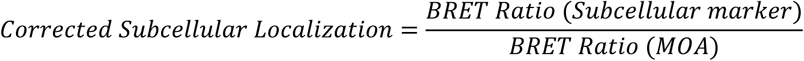

Total receptor expression was quantified from the mean donor luminescence values.

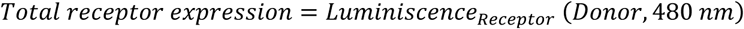

All corrected datasets were analyzed statistically using ordinary one- or two-way ANOVA followed by Tukey’s multiple-comparisons test.

### SNAP-tag-based detection of cell surface expression

Detection of cell-surface receptor was performed by SNAP-based labeling with a terbium fluorophore probe (SNAP-Lumin4-Tb; Revvity, Inc., Cat#SSNPTBD) in the extracellular milieu^58^. HEK293A cells were seeded at a density of 20,000 cells per well into poly-D-lysine-coated white 384-well plates and co-transfected with 10 ng of N-terminally FLAG-SNAP-tagged GPCR plasmid together with 30 ng of pcDNA3.1, giving a total of 40 ng DNA per well. Transfections were performed using Lipofectamine™ 2000 (Thermo Fisher Scientific, Cat#11668027) at a DNA-to-reagent ratio of 1:2 (w/v). Twenty-four hours later, the culture medium was replaced with Opti-MEM™ I Reduced Serum Medium containing 100 nM SNAP-Lumi4-Tb, and cells were incubated for 1 hour at 4 °C. Following labeling, cells were washed six times with an internalization buffer composed of Hanks’ Balanced Salt Solution (HBSS), 1 mM CaCl₂, 1 mM MgCl₂, 20 mM HEPES, and 0.1% (v/v) bovine serum albumin (BSA) at pH 7.4. Cell surface receptor expression was then determined by adding internalization buffer to each well and measuring the SNAP-Lumi4-Tb fluorescence signal at 620 nm. Statistical differences between groups were evaluated using one-way ANOVA followed by Tukey’s multiple-comparison post hoc test.

### Expression and purification of CXCR4

Based on previous protocols^30^, human wildtype CXCR4 (residues 2–362), N^3.35^119S, W^4.50^161Y, and L150^4.40^C/Q233^6.29^C mutants were heterologously expressed in Sf9 insect cells using the Bac-to-Bac baculovirus expression system (Thermo Fisher Scientific). For this, their cDNA was cloned into a pFastBac1 vector incorporating an N-terminal polyhedrin (PolH) promoter and hemagglutinin (HA) signal peptide, together with C-terminal His₁₀, FLAG and AviTag™ affinity tags. Cells expressing the CXCR4 constructs were harvested 48 h after infection, and membrane fractions were prepared by Dounce homogenization in hypotonic buffer, with repeated high-salt washing steps performed for CXCR4 membranes. Prior to detergent extraction, CXCR4 membranes were treated with 100 µM IT1t dihydrochloride (MedChemExpress, Cat#HY-101458A). Membrane proteins were solubilized for 1 h in 50 mM HEPES (pH 7.5), 400 mM NaCl, and 0.75% (w/v) n-dodecyl-β-D-maltopyranoside (DDM; Anatrace, Cat#D310) without addition of cholesteryl hemisuccinate (CHS), with EDTA-free protease inhibitor cocktail (Sigma-Aldrich, Cat#5056489001) included during CXCR4 purification. Insoluble material was removed by centrifugation (50,000 × *g*, 30 min), and clarified lysates were supplemented with 20 mM imidazole; Recombinant receptors were captured by metal affinity chromatography using TALON® Superflow™ resin (Cytiva, Cat#28-9575-02) through overnight binding at 4 °C in the presence of 1 U mL⁻¹ DNase I (Thermo Scientific, Cat#90083). The following day, the beads were washed with detergent-containing buffers of progressively reduced DDM concentrations. Proteins were eluted with 500 mM imidazole, desalted using PD MiniTrap G-25 columns (Cytiva, Cat#28918007), and exchanged into 50 mM HEPES (pH 7.5), 400 mM NaCl, and 0.025% (w/v) DDM. Samples were concentrated using 100-kDa molecular weight cut-off protein concentrators (Thermo Fisher Scientific, 88533), and protein purity was verified by SDS-PAGE. CXCR4 concentration was determined spectrophotometrically at 280 nm using the calculated extinction coefficients (65,820 M^−1^ cm^−1^ for WT and N^3.35^119S CXCR4, 61,810 for W^4.50^161Y CXCR4, and 65,320 for L150^4.40^C/Q233^6.29^C CXCR4), after which purified proteins were snap-frozen in liquid nitrogen and stored at −80 °C until further use.

### Nanodisc reconstitution and CXCR4 biotinylation

Nanodiscs were assembled using phospholipids obtained from Sigma-Aldrich using an adapted protocol of^28^. 1-palmitoyl-2-oleoyl-*sn*-glycero-3-phosphocholine (POPC) and 3β-hydroxy-5-cholestene (cholesterol; Merck, Cat#C3045), initially dissolved in chloroform, were combined at two molar compositions (100% POPC, or 4:1 POPC:cholesterol), dried under a gentle stream of nitrogen, and subsequently resuspended to a final lipid concentration of 100 mM in cholate-containing buffer (50 mM HEPES pH 7.5, 150 mM NaCl, supplemented with sodium cholate at a final detergent concentration of 24 mM). To favor the incorporation of single receptor molecules per nanodisc, the MSP1D1 scaffold protein was used, generating nanodiscs with an approximate diameter of 10 nm^59^. Nanodiscs were reconstituted by mixing purified CXCR4 constructs, untagged MSP1D1 scaffold protein, and lipids at a molar ratio of 0.1:1:76 (CXCR4:MSP1D1:lipid), while maintaining a final cholate concentration above 20 mM. For empty nanodiscs, receptor protein was omitted and replaced with buffer. Following incubation at RT for 20–30 min with gentle agitation, detergent removal was initiated by adding Bio-Beads SM-2 resin (Bio-Rad, Cat#152-3920), allowing overnight nanodisc self-assembly at 4 °C. After removal of the resin, the CXCR4 samples used for smFRET were biotinylated using BirA. For this, CXCR4 nanodiscs were incubated with 0.0125 mg/ml BirA, 0.2 mM biotin, 10 mM ATX and 10 mM MgCl_2_ for 2 h at RT without shaking. Nanodisc preparations were then purified by size-exclusion chromatography using a ProteoSEC Dynamic 11/30 6-600 HR column (Protein Ark/Bionordica, Cat#SEC-D-11/30-6-600) equilibrated in HEPES buffer (25 mM HEPES pH 7.5, 50 mM NaCl). Fractions containing nanodiscs were pooled and assessed by SDS-PAGE where appropriate. Purified samples were concentrated to approximately 1–30 μM and stored either at 4 °C for short-term use or at −80 °C for extended preservation.

### *In vitro* mini-Gα_o_ binding to CXCR4

*In vitro* G protein binding was examined by a Fluorescence saturation binding assay (FSBA) based on^60^. Purified His_10_-tagged WT CXCR4, N119S, or W161Y was reconstituted in nanodiscs containing untagged MSP1D1 with POPC:cholesterol (4:1 molar ratio). For empty nanodiscs used as a control for non-specific binding, His_8_-tagged MSP1D1 was reconstituted without receptor in the same lipid composition. TALON® Superflow™ beads (10 µl of resin per tested condition) were transferred to a new tube, where they were centrifuged at 100 × g for five minutes, after which the beads were washed in nanodisc buffer (25 mM HEPES pH 7.5, 50 mM NaCl, supplemented with 0.1% BSA, w/v). The wash was repeated a second time, after which the supernatant was removed, and 0.5 ng was added per 10 µl of resin. The nanodiscs were allowed to bind to the beads for at least 30 min at 4 °C in a rotating disk. Upon incubation, the samples were washed twice, a final concentration of 5µM GFP-mini-Gα_o_ was added and mixed, which was then divided over PCR tubes (equivalent amount for 10 µl of resin per tube). Nanodisc buffer, 0.5 µM CXCL12, or 0.5 µM IT1t was added to each tube and allowed to incubate at RT for at least 30 min. The samples were then kept at 4 °C to prevent fast dissociation; they were spun down, the supernatant was removed using a multi-channel pipette, and nanodisc buffer was added. This process was repeated twice, after which the beads were incubated with nanodisc buffer supplemented with 300 mM imidazole, and bead-bound nanodiscs were allowed to dissociate for at least 15 min. The supernatants were then collected, and fluorescence was measured at 509 nm using a CLARIOstar microplate reader (BMG Labtech).

### smFRET sample preparation

For single-molecule FRET (smFRET) experiments, 2 nmol of receptor was thawed and incubated overnight at 4 °C under continuous rotation with 14 nmol each of Alexa Fluor™ 555 C2 maleimide (AF555; molecular weight 1250 g/mol; Thermo Fisher Scientific, Cat#A20346) and Cy®5 maleimide (Cy5; molecular weight 641.25 g/mol; Sigma-Aldrich/Merck, Cat#GEPA15131). Following incubation, excess unbound fluorophore was removed by repeated dilution and concentration using a 100 kDa molecular weight cut-off centrifugal filter (Amicon Ultra 100 kDa centrifugal filters; MilliporeSigma, Cat#UFC510024), and the sample volume was reduced to approximately 100 µL. Labeling efficiency was determined by measuring absorbance at 280 nm, 555 nm, and 645 nm using extinction coefficients of 65,320 M⁻¹cm⁻¹ for the receptor, 150,000 M⁻¹cm⁻¹ for AF555, and 250,000 M⁻¹cm⁻¹ for Cy5, respectively. Prior to calculating receptor concentration, the absorbance contribution of the attached fluorophores at 280 nm was subtracted. The entire labeled receptor preparation was subsequently incorporated into untagged MSP1D1 nanodiscs according to the nanodisc reconstitution protocol described in the section "Nanodisc reconstitution and CXCR4 biotinylation."

### smFRET data collection and analysis

Single-molecule fluorescence measurements were performed using a custom-built prism-based TIRF imaging system, as previously described. An Olympus IX73 inverted microscope equipped with a 60x water-immersion objective (Olympus, 1.49 NA) was used for data collection^61,62^. Before the image acquisition, quartz slides (Technical Glass Products, Inc., Painesville, OH, USA) and coverslips were passivated with polyethylene glycol (m-PEG-SVA) and 3% (w/w) biotin-PEG-SVA (Laysan Bio Inc., Arab, AL, USA) to suppress non-specific adsorption^63,64^. A sample chamber was prepared using double-sided tape, grease, and a coverslip. The chamber was first passivated with 0.2 mg/ml of NeutrAvidin (Thermo Scientific, Cat#PI3100) and then washed with imaging buffer A (25 mM HEPES, pH 7.5, 150 mM NaCl, 1 mM propyl gallate, 2 mM Trolox, Thermo Scientific Cat#AC218940050) as described previously^30^. A doubly fluorescent-labeled and biotin-conjugated CXCR4 nanodisc sample was then diluted in buffer A, injected into the slide chamber, and incubated for 10 minutes. The samples were then washed with buffer A, followed by an imaging buffer containing an oxygen scavenger system (OSS) with 2 mM Trolox, 5 mM protocatechuic acid (PCA, Sigma-Aldrich 37580-25G-F), and 3 U/ml protocatechuate-3,4-dioxygenase (rPCO) (Oriental Yeast, Cat#46852004). The sample was illuminated with a 532 nm laser, and fluorescence emission from donor and acceptor fluorophores was collected on an EMCCD camera (Andor Technology) at room temperature with an exposure time of 100 ms. Images were recorded using a custom single-molecule data acquisition program^65^. The data collected was processed using custom scripts written in IDL (Interactive Data Language, Harris Geospatial Solutions, Inc.). The source code used for data acquisition and extraction is available in a repository at https://github.com/Ha-SingleMoleculeLab. Experiments were repeated for 2.5 µM CXCL12 and the 1 µM small molecule IT1t. The data presented represent the combined results of at least three separate experiments for each condition.

We used custom-built scripts to generate FRET traces as described previously (https://github.com/rpauszek/smtirf) ^65^. Fluorescence intensity trajectories for the donor and acceptor channels were first corrected for donor spectral bleed-through and background fluorescence signal. Apparent FRET efficiency at each time point was then calculated using the expression *E_app_* = *I*_A_/(*I*_A_+*I*_D_), where *I*_D_ and *I*_A_ denote the background-corrected donor and acceptor fluorescence intensities, respectively. Individual fluorescence trajectories were subsequently manually inspected for single-step photobleaching events for both fluorophores and to verify the expected anticorrelated donor–acceptor intensity changes. These criteria were used to confirm that each fluorescent spot corresponded to a single receptor molecule labeled with one donor and one acceptor fluorophore. Multiple single-molecule FRET trajectories were used to generate single-molecule FRET histograms for each condition tested. FRET trajectories were pooled and analyzed separately using STaSI to identify the corresponding conformational states^66^. FRET histograms were generated and fitted using Igor Pro software (Wavemetrics, Lake Oswego, OR, USA). Individual fluorescence trajectories were analyzed using a Hidden Markov Model (HMM). Transition density plots (TDPs) illustrating the connectivity between individual FRET states were subsequently generated as previously described ^62,67^.

## Data analysis

Dose–response curves were analyzed by fitting a four-parameter nonlinear regression model (log[agonist] versus response) with a variable slope. To minimize variability between experimental days, data from independent experiments were compared using ordinary two-way ANOVA assessing main column effects, followed by Dunnett’s multiple-comparison test. Statistical analyses were performed using GraphPad Prism version 10.6.

## Data availability

All data are available upon request to.

## Acknowledgements

Research in the Gustavsson lab was supported by Villum Fonden (grant 00025326), Carlsberg Foundation (grant CF19-0320) and Independent Research Fund Denmark (grant 3103-00230B). Research in the Lamichhane lab was supported by the National Institutes of Health (grant R35GM142946).

## Author information

These authors contributed equally: Jose Fernández-González, Carolina Ferrera-Mena

## Authors and Affiliations

**Department of Biomedical Sciences, University of Copenhagen, Copenhagen, Denmark**

Fernando Salgado-Polo, Jose Fernández-González, Carolina Ferrera-Mena, Philip Ben Rainsford, Rajesh Regmi, Martin Gustavsson

**Department of Biochemistry & Cellular and Molecular Biology, University of Tennessee, Knoxville, Tennessee, USA**

Suresh Subedi, Sriram Tiruvadi Krishnan, Rajan Lamichhane

## Contributions

F.S.-P, M.G., and R.L. were responsible for study conceptualization and experimental design. F.S.-P, J.F.-G, C.F.-M, S.S., S.T.K., P.B.R., and R.R. conducted wet-lab experiments, data analysis, and visualization. F.S.-P wrote the original draft, which was revised and polished by R.L. and M.G.

## Corresponding author

Correspondence to Martin Gustavsson.

