## Extended Data Figures 1-5 and Table 1 for "Chemokine receptor activity is differentially regulated by membrane cholesterol"

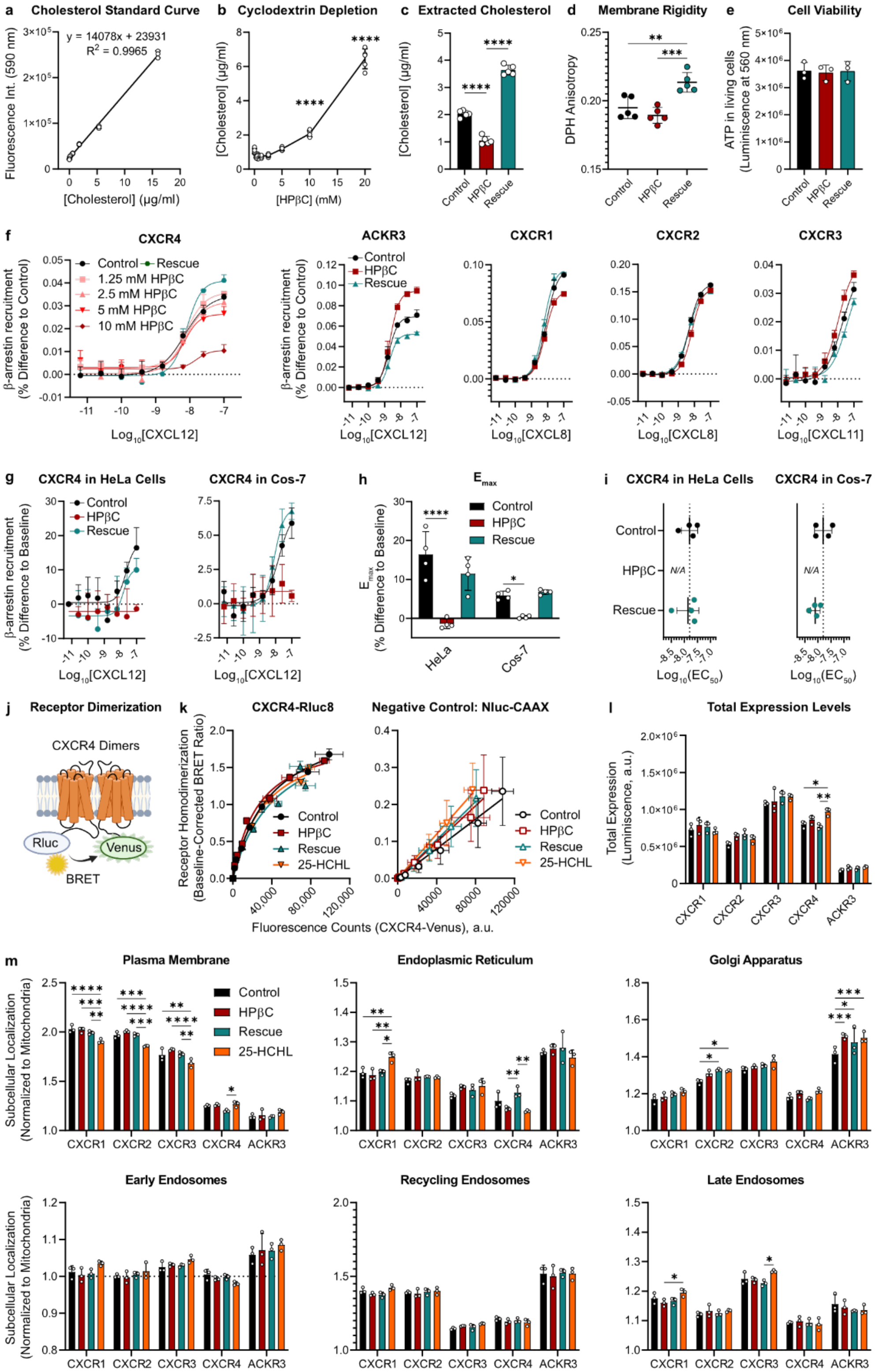


**Extended Data Fig. 1: Cholesterol selectively modulates β-arrestin recruitment and internalization of chemokine receptors.** (Legend continues in the next page).

**Extended Data Fig. 1: Cholesterol selectively modulates β-arrestin recruitment and internalization of chemokine receptors.** **a**,**b**, Linear regression of the fluorescence levels measured upon oxidation of known concentrations of cholesterol (**a**). The resulting function was used to calculate the concentration of cholesterol that was extracted from HEK293A cells (**b**). **c**, Amount of cholesterol extracted from control, HPβC-treated, and cholesterol-rescued HEK293A cells. **d**, The fluorescent, membrane-embedded probe DPH was used to detect changes in rigidity in the membrane of HEK293A cells upon cholesterol extraction or rescue. Increased anisotropy levels related to increased membrane rigidity. **e**, Effects of the treatments with HPβC on HEK293A cell viability, measured with CellTiter-Glo® Luminescent Cell Viability Assay. **f**, Dose-response curves for β–arrestin2 recruitment corrected to the baseline of the control group. Data refers to **Fig.1c**,**e**, where each curve was corrected to its own baseline. **g**, Effect of decreased cholesterol on the recruitment of β–arrestin2 to CXCR4 in HeLa (left) and Cos-7 (right) cells. **h**,**i**, Changes in the efficacy (**h**) and the potency (**i**) of β–arrestin2 to CXCR4 in HeLa and Cos-7 cells. **j**, Schematic representation of the assay used to study the role of cholesterol in chemokine receptor homodimerization; created with BioRender.com. **k**, Effect of decreased cholesterol on the constitutive homodimerization of CXCR4 in HEK293A cells. Cells were transiently transfected with C-terminally Rluc8- or Venus-tagged CXCR4, which resulted in higher BRET levels upon dimerization. Data was fitted by a non-linear regression to a hyperbola, whose shape is indicative of specific binding events. Membrane-anchored Nluc-CAAX was used as a control for lack of binding. **l**, Total expression levels of the chemokine receptors shown in **Fig.1** in transiently transfected HEK293A cells. Luminescence was collected from non-stimulated cells expressing C-terminally Rluc3-tagged constructs. **m**, Effect of decreased cholesterol and rescue on the co–localization of CXCR1, CXCR2, CXCR3, CXCR4 and ACKR3 with the indicated Venus–tagged subcellular localization markers. Plasma membrane levels refer to the data shown in **Fig.1i-j**. N = 5 (**a-d**) or N = 3 (**e-i**,**k-m**) independent experiments, performed in technical triplicates. All data are shown as mean ± SD. The asterisk symbols indicate statistically significant differences (*p < 0.05, **p < 0.01, ***p < 0.001, ****p < 0.0001) between the indicated groups, determined by one-way (**b**,**c**,**d**,**e**,**h**,**i**) or two-way (**l**,**m**) ANOVA. Only significant differences are shown.

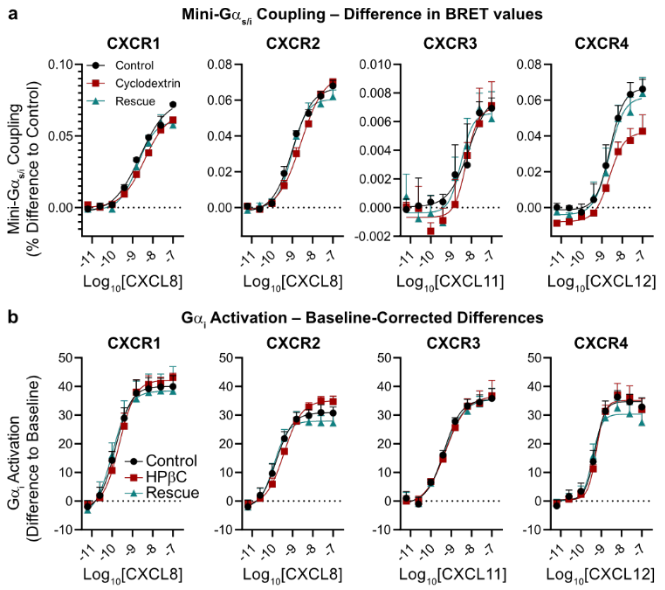


**Extended Data Fig. 2: Cholesterol modulates constitutive and ligand-induced G protein signaling in chemokine receptors**. **a**, Dose-response curves for mini-$G\alpha_{s/i}$ coupling corrected to the baseline of the control group. Data refers to **Fig.2b**, where each curve was corrected to its own baseline. **b**, Dose-response curves for $G\alpha_i$ activation, where each group was corrected to its own baseline. Data refers to **Fig.2f**, where all curves were corrected the baseline of the control group. In all panels, N = 3 independent experiments, performed in technical triplicates. All data are shown as mean ± SD.

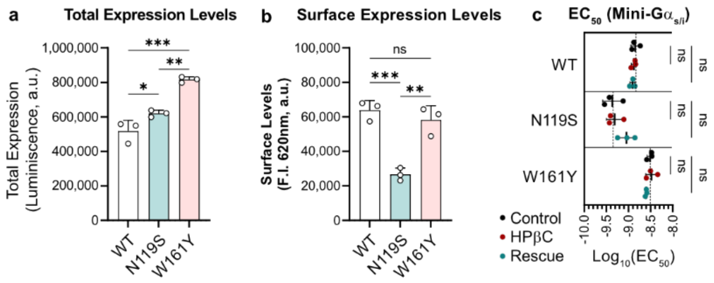


**Extended Data Fig. 3: Cholesterol facilitates the activation of CXCR4, enabling efficient G protein coupling**. **a**, Total expression levels of CXCR4 and its mutants in transiently transfected HEK293A cells. Luminescence was collected from non-stimulated cells expressing C-terminally Rluc3-tagged constructs. **b**, Surface expression levels of N-terminally SNAP-tagged CXCR4 constructs labeled with SNAP-Lumi4®-Tb at 4 °C. **c**, Effect of cholesterol extraction and rescue on the potency for mini-G$\alpha_{s/i}$ coupling to WT, N119S, and W161Y CXCR4 constructs. Data refers to the dose-response curves shown in **Fig.3d**. N = 3 independent experiments, performed in technical triplicates. All data are shown as mean ± SD. The asterisk symbols indicate statistically significant differences (**$p < 0.01$, ***$p < 0.001$, ****$p < 0.0001$, ns denotes non-significant changes) between the indicated groups, determined by one-way ANOVA.

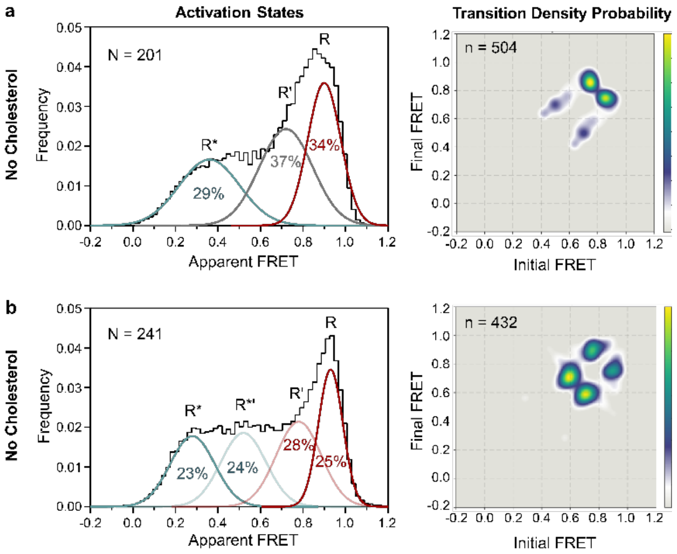


**Extended Data Fig. 4: Cholesterol facilitates the activation of CXCR4, enabling efficient G protein coupling**. **a-b**, Changes in the activation states of IT1t-bound in the absence (**a**) and presence (**b**) of 20% cholesterol using single–molecule fluorescence resonance energy transfer (smFRET). Left panels show histograms of single-molecule FRET states, while the right panels show two-dimensional transition density probability (TDP) plots for the transitions between FRET states. N represents the number of single-molecule traces used to generate histograms, and n represents the total number of transitions across the states.

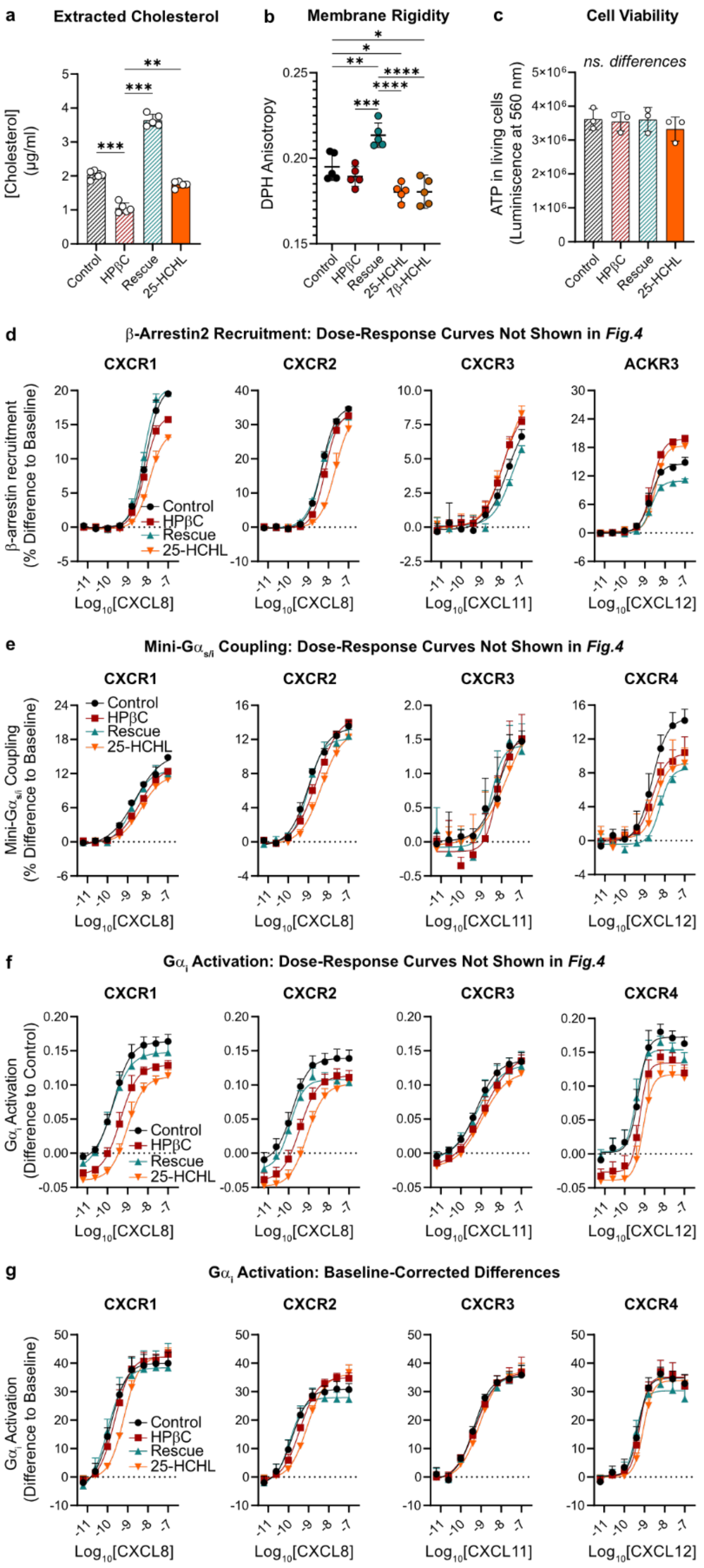


**Extended Data Fig. 5: Oxysterols fail to rescue the effects of cholesterol on chemokine receptor activity**. (Legend continues in the next page).

**Extended Data Fig. 5: Oxysterols fail to rescue the effects of cholesterol on chemokine receptor activity**. **a**, Amount of cholesterol extracted from control, HPβC-treated, cholesterol-rescued, and 25-HCHL-rescued HEK293A cells. **b**, Changes in membrane rigidity as a result of decreasing the amount of cholesterol or rescuing the cells with (hydroxy)cholesterol-loaded HPβC. Increase in the anisotropy levels of the fluorescent, membrane-embedded probe DPH accounted for increases in rigidity in the cells' membrane. **c**, Effects of the treatments with HPβC on HEK293A cell viability, measured with CellTiter-Glo® Luminescent Cell Viability Assay. **d**, Dose-response curves for β–arrestin2 recruitment, where each curve was corrected to its own baseline. Data refers to the quantifications shown in **Fig.5c**. **e**, Dose-response curves for mini-$G\alpha_{s/i}$ coupling, where each curve was corrected to its own baseline. Data refers to the quantifications shown in **Fig.5d**. **f**,**g**, Dose-response curves for $G\alpha_i$ activation, where each group was corrected to the baseline of the control curve (**e**) or to its own baseline (**f**). Data refers to the quantifications shown in **Fig.5e**. (**a-c**) The asterisk symbols indicate statistically significant differences (**$p < 0.01$, ***$p < 0.001$, ns denotes non-significant changes) between the indicated groups, determined by one-way ANOVA. N = 5 (**a-b**) or N = 3 (**c-f**) independent experiments, performed in technical triplicates. All data are shown as mean ± SD. (**a**) Only significant differences are shown.

**Extended Data Table 1: FRET efficiencies for each CXCR4 conformational state.**

| ***FRET Efficiency ($E_{app}$)*** | **Active (R*)** | **Active-like (R*')** | **Inactive-like (R')** | **Inactive (R)** |
|---|---|---|---|---|
| *Apo, no CHL* | 0.31 | | 0.72 | 0.90 |
| *Apo, 20% CHL* | 0.25 | 0.52 | 0.85 | 0.94 |
| *CXCL12, no CHL* | 0.25 | 0.52 | | 0.90 |
| *CXCL12+, 20% CHL* | 0.25 | 0.46 | 0.72 | 0.93 |
| *IT1t, no CHL* | 0.29 | | 0.72 | 0.91 |
| *IT1t, 20% CHL* | 0.28 | 0.52 | 0.78 | 0.93 |
